# Environmental influences on acoustic divergence in *Rhinolophus* bats of the Western Ghats-Sri Lanka region

**DOI:** 10.1101/661314

**Authors:** Kadambari Deshpande, Nachiket Kelkar

**Author notes:** Corresponding author: Kadambari Deshpande Present address: PhD student, Academy of Conservation Science and Sustainability Studies, Ashoka Trust for Research in Ecology and the Environment (ATREE), Royal Enclave, Srirampura, Jakkur Post, Bangalore 560064, Karnataka, India.

## Abstract

According to the acoustic adaptation hypothesis, environmental and biogeographic factors such as atmospheric humidity can influence divergence of acoustic signals and speciation in high duty-cycle echolocating bats (e.g. *Rhinolophus* sp.), although this remains disputed. In this study we tested the hypothesis that Resting Frequency (RF) would decrease with increasing humidity along a large latitudinal gradient (6**°**-21**°** N), for four *Rhinolophus* species with different evolutionary histories, in the Western Ghats-Sri Lanka (WGSL) region. We conducted acoustic recordings and compiled published information on RFs of stationary *Rhinolophus indorouxi, R. rouxi, R. beddomei*, and *R. lepidus* from 40 roosts in 18 localities of the WGSL. These data comprised of recordings made with different devices and with different settings. Hence, due to the unknown measurement error involved in the recorded RFs, it was not possible to conduct conventional regression analyses to test our hypotheses. Hence, we qualitatively assessed effects of Relative Humidity (RH) and other environmental variables by interpreting only the sign, but not the magnitude of the RF responses (from the slopes of generalized least squares regression models). We also tested how RF and RH varied across biogeographic zones, and with bat body size. RFs of the Miocene-diverged species *R. indorouxi* and *R. rouxi* were higher at lower RH, as expected. In contrast, RF of the Pleistocene-diverged species *R. beddomei* and *R. lepidus* were higher at higher RH. Elevation and rainfall also emerged as important predictors of RF variation in these species. Bat body size differed in dry and humid regions of the WGSL. RF variation was not consistent across biogeographic zones. The cryptic, phonically differentiated sibling species *R. indorouxi* and *R. rouxi* co-occurred only in mid-elevation zones along the Western Ghats escarpment. The variable but significant influences of humidity and correlated factors on RF suggest the importance of environmentally mediated acoustic divergence in different *Rhinolophus* species in the WGSL. We propose some hypotheses on interacting effects of environmental and phylogenetic factors on acoustic divergence in *Rhinolophus* bats of the WGSL. These ideas could be further tested with phylogenetic and acoustic studies, as more consistent and comparable data on these species become available in the future.

## Introduction

Trait divergence in animals is an important trigger for ecological speciation (Rundle and Nosil, 2005; Schluter, 2001; Wilkins et al., 2013). Adaptive processes such as ecological and sexual selection, or neutral, non-adaptive processes such as genetic drift can produce divergence (Fitzpatrick et al., 2009; Wilkins et al., 2013). Among adaptive processes, climatic and environmental selection pressures can influence rapid divergence of specialized and functionally important traits such as acoustic signals (Jones and Teeling, 2006; Wilkins et al., 2013). Phonic differentiation in ultrasonic echolocation calls is known in many insectivorous bat populations, making them ideal systems to test the effects of environmental variability and change on acoustic divergence (Jones, 1997; Sedlock and Weyandt, 2009; Stoffberg et al., 2011). In high duty-cycle echolocating bats, e.g. horseshoe and leaf-nosed bat species, acoustic frequencies are held constant over longer pulse durations with the use of Doppler Shift Compensation to match the sensitivity of acoustic fovea (Clare et al., 2013; Jacobs and Bastian, 2018; Schuchmann and Siemers, 2010; Stoffberg et al., 2011). This is to resolve minute differences in hearing for spectral partitioning of signals from the echoes (Bastian and Jacobs, 2015; Heller and von Helversen, 1989; Kingston et al., 2000; Raw et al., 2018; Roberts, 1972), to aid effective prey capture against highly cluttered backgrounds (Clare et al., 2013; Bastian and Jacobs, 2015). Further, *Rhinolophus* bats can also discriminate and recognize the sex of conspecific members from their populations from other conspecific or heterospecifics (Lin et al., 2016; Raw et al., 2018; Schuchmann et al., 2012). Such extreme specialization in high duty-cycle echolocation could have also led to phenotypic convergences in *Rhinolophus* bat species, even irrespective of phylogenetic relatedness (Jacobs et al., 2013, 2016; Jacobs and Bastian 2018). High specialization is indicated by the limited diversity of foraging niches used by bats in this genus (Jacobs et al., 2016). But cryptic speciation through acoustic differentiation is widespread in horseshoe and leaf-nosed bats (Chiroptera: Rhinolophidae, Hipposideridae), as seen across tropical and subtropical regions in South Africa, Asia, China, southern Europe, and Australasia (e.g. Chen et al., 2009; Guillen et al., 2000; Jiang et al., 2013, 2010a; Odendaal et al., 2014; Srinivasulu et al., 2019; Sun et al., 2013; Xu et al., 2008).

Jones (1997), Jones and Teeling (2006), and Jacobs et al. (2017) emphasized the relative importance of ecological processes in shaping acoustic divergence, i.e. “sensory drive” in high duty-cycle bats, over and above phylogenetic constraints or neutral processes like genetic drift. Numerous studies have supported stronger effects of non-neutral, environment or climate-driven selection pressures than drift on acoustic divergence (Jacobs et al., 2017; Jacobs & Mutumi, 2018; Mutumi et al., 2016). Mutumi et al. (2017) distinguished effects of environmentally mediated selection from drift by showing that variation in *Rhinolophus* bat phenotypic traits and frequencies did not correlate with geographic distance. Adaptive plasticity in response to environmental heterogeneity, coupled with geographic or genetic isolation, can also affect acoustic divergence both in sympatric and allopatric species (Maluleke et al., 2017; Odendaal et al., 2014; Russo et al., 2007; Stoffberg et al., 2012, 2011; Yoshino et al., 2008). Acoustic divergence could be driven also by social or sexual selection (Kingston et al., 2001; Mallet et al., 2009; Puechmaille et al., 2014), by specialization of signals for communication (Thabah et al. 2006), cultural drift (Yoshino et al., 2008; Lin et al., 2014, Xie et al., 2017), interspecific competition for prey (Schuchmann and Siemers, 2010), or resource availability (Kelly, 2008) possibly in relation to environmental variation.

Among the numerous studies reporting environmentally driven variation in acoustic frequency, atmospheric humidity has received the most attention (Chen et al. 2009; Flanders et al. 2011; Guillen et al., 2000; Jiang et al. 2013, 2010a, 2010b; Sun et al. 2013; Xu et al. 2008). As per the sensory drive framework (that includes the acoustic adaptation hypothesis: Sun et al., 2013; Wilkins et al., 2013), greater atmospheric humidity is thought to alter *Rhinolophus* bat frequencies (Jiang et al., 2015), as it can directly affect signal transmission, sound absorption, and echo reception (Goerlitz, 2018; Snell-Rood, 2012). However, others have disputed the influence of humidity in driving frequency attenuation, showing that acoustic transmission at fine scales remains unaffected despite changes in humidity up to 10% (e.g. Armstrong and Kerry, 2011; Goerlitz, 2018). Humidity could also be linked to habitat preferences of bats or insect prey size (Guillen et al., 2000; Jiang et al., 2010b), thus indirectly altering bat frequencies. The James’ rule (James, 1970) predicts larger body sizes to be favoured in cool, dry areas, whereas moderate-sized or small individuals of the same species to be favoured in hot, humid areas due to metabolic constraints. Body sizes varying in response to climatic factors (Jacobs and Bastian 2018) can cause acoustic divergence in relation to allometry. Stoffberg et al. (2011) also provided support for the evolution of adaptive complexes between body size and echolocation in *Rhinolophus*. To understand modes of acoustic divergence in *Rhinolophus* bats, we thus need to understand linkages between resting frequencies, body size, environmental factors such as humidity, and evolutionary histories across large-scale climatic gradients (Chen et al., 2009; Jones, 1997; Mao et al., 2010; Mutumi et al., 2017; Odendaal et al., 2014; Stoffberg et al., 2011).

The Western Ghats-Sri Lanka (WGSL) region offers an excellent setting to test hypotheses on divergence. The WGSL stretches over 15 degrees of latitude (6**°**-21**°** N) spanning a wide range of environmental heterogeneity (humidity, elevation, rainfall, topography and vegetation types; Bose et al. 2015). Orographically, the region has five potential biogeographic barriers due to physical discontinuities or gaps. The effects of these barriers, e.g. the Palghat Gap, on divergent selection and speciation in birds, anurans, plants etc. have been well documented (Bose et al., 2015; Gunawardene et al. 2007; Robin et al., 2010; Vijaykumar et al., 2016). Robin et al. (2011) have reported congruence of song type and genetic variation in short-wing (songbird species) populations of the Western Ghats. Environmental and biogeographic influences on acoustic divergence in bats have not yet been assessed in the WGSL. However, their effects on cryptic speciation are evident from the recent genetic description of two phonic types of the rufous horseshoe bat *Rhinolophus rouxi*, later assigned to two cryptic sibling species (Chattopadhyay et al., 2012). The occurrence of four species of *Rhinolophus* bats with distinct evolutionary histories allows for interesting comparisons of acoustic variation related to environmental effects.

In this paper, we report results of our preliminary investigations on the environmental and biogeographic correlates of acoustic divergence in four *Rhinolophus* bat species in the Western Ghats-Sri Lanka (WGSL) region. Specifically, we tested the “sensory drive” prediction of the acoustic adaptation hypothesis (Wilkins et al., 2013) that resting frequency will reduce as atmospheric humidity increases, as against the null hypothesis of no relationship with humidity. We also tested the effects of other environmental variables such as elevation, rainfall, and vegetation on variation in Resting Frequencies (RFs) of these bats. Analyses were done separately for the four species to account for their phylogeographic distinctiveness (Ith et al., 2016). We first assessed variation in RF along latitudinal climatic gradients (Jiang et al., 2015) and across biogeographic zones of the WGSL. Then we checked the correlation of body size with RF (Stoffberg et al., 2011) and humidity (Malukele et al., 2017; Mutumi et al., 2016) as a test of the James’ rule. This helped us test if humidity influenced frequency in relation to body size or not (Mutumi et al., 2016; Russo et al., 2007; Stoffberg et al., 2012). Based on our results, we discuss how environmental factors may have influenced divergence in four *Rhinolophus* species across the WGSL. Our aim is to suggest potential hypotheses that could be tested with rigorous studies in the future.

## Methods

### Study area

The Western Ghats and Sri Lanka (WGSL) biodiversity hotspot has high floral and faunal endemism (Gunawardene et al. 2007). The WG in India run roughly parallel to the western coastline of India for c.1600 km, in the NNW-SSE direction, and are geologically continuous with the central highlands of Sri Lanka (Gunawardene et al., 2007; Fig. 1). The island of Sri Lanka is separated from the Indian peninsula by 35-40 km, but intermittent connections and species exchanges have occurred historically and through the Quaternary period (Bose et al., 2015). The WG edge is a shoulder-type escarpment straddling the western boundary of the relict mountain chain (Gunnell, 1997; Harbor and Gunnell, 2007). Elevation highs or “massifs” of the Nilgiris, Anamalais, and Palnis along the WG escarpment affect rainfall and humidity regimes along the WGSL (Vijayakumar et al., 2016). Rainfall intensity and duration reduce, and rainfall becomes more seasonally pulsed, as one travels northward (Gunnell, 1997; Krishna Prasad et al., 2008). The Palghat Gap within the WG and the gap between WG-Sri Lanka are the two main biogeographic barriers that have driven endemic radiations of many taxa (e.g. Gunawardene et al., 2007; Robin et al., 2010; Vijayakumar et al., 2016). From north to south, the Goa Gap, Cauvery valley, and the Palghat and Shenkottah Gaps also form distinctive biogeographic zones (Purushotham and Robin, 2016; Ramachandran et al., 2017; Robin et al., 2010). These zones roughly correspond with latitudinal gradients in humidity, rainfall seasonality, and elevation (Bose et al., 2015). Figure 1 describes the delineation of these zones in the WGSL. Twelve bat species from the WGSL (Bates and Harrison, 1997; Csorba et al., 2003) belong to the high duty-cycle bat families Rhinolophidae (5 *Rhinolophus* spp.) and Hipposideridae (7 *Hipposideros* spp.), which also occur in other areas of peninsular India (Bates and Harrison, 1997).

**Figure 1.**
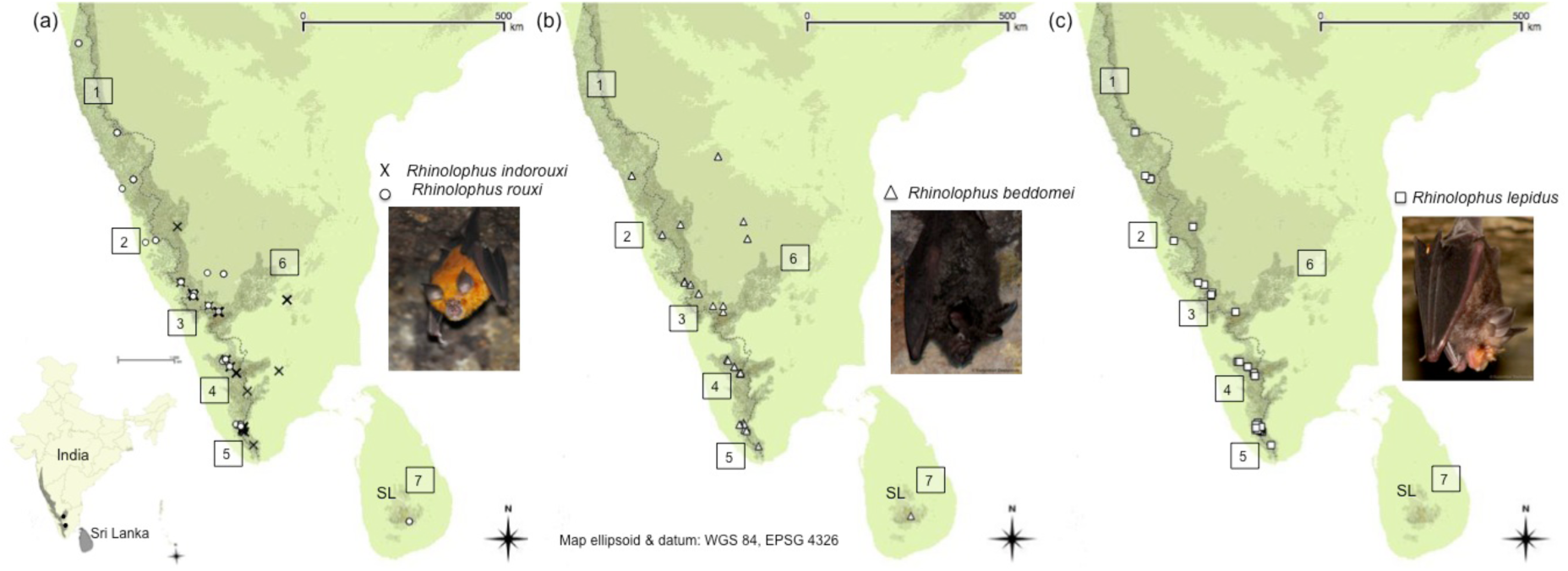
Locations of recorded occurrence of the four *Rhinolophus* species in the Western Ghats and Sri Lanka region, from which resting frequencies were selected for analyses. Numbers represent different biogeographic zones as follows: 1) North of Goa Gap (N.G), 2) Goa Gap to Cauvery valley (G-CV), 3) Cauvery valley to Palghat Gap (CV-PG), 4) Palghat Gap to Shenkottah Gap (PG-SG), 5) South of Shenkottah Gap (S.SG), 6) Peninsular India and Eastern Ghats region (P.EG) for comparison, and 7) Sri Lanka (SL). (a) *Rhinolophus rouxi* and *R. indorouxi* are cryptic sibling species (Chattopadhyay et al., 2012) forming a possible *R. rouxi* ‘complex’ (Thomas, 1997; Wordley et al., 2014). (b) *R. beddomei* calls were not available from North of Goa, where it occurs (Bates and Harrison, 1997). (c) *R. lepidus* is likely absent from Sri Lanka (Csorba et al., 2003).

### Species identification and taxonomic assignments

*Rhinolophus* species known from the WGSL are: 1) *R. rouxi* Temminck, 1835, 2) *R. beddomei* Andersen 1905, 3) *R. lepidus* Blyth, 1844, 4) *R. pusillus* Temminck, 1834, (Bates and Harrison, 1997; Csorba et al., 2003), and 5) *R. indorouxi*, a sibling species of *R. rouxi* recently described by Chattopadhyay et al. (2012). For the study we focused on four species (*R. rouxi, R. beddomei, R. lepidus*, and *R. indorouxi*). We never encountered *R. pusillus* in our surveys. We standardized species assignments by following these taxonomic criteria: 1) Sri Lankan records of *R. rouxi* were referred to the subspecies *R. rouxi rubidus*, and of *R. beddomei* to *R. beddomei sobrinus*, 2) past records of *R. luctus*, subject to taxonomic uncertainty, were assigned to *R. beddomei*, following Csorba et al. (2003), and 3) *R.* cf. *indorouxi* was called *R. indorouxi*, following Chattopadhyay et al. (2012). Phylogenetic and evolutionary information from Csorba et al. (2003) and Chattopadhyay et al. (2012) showed that *R. rouxi* evolved from *R. indorouxi* during Miocene drying. *R. beddomei* and *R. lepidus* likely colonized the WGSL in the Pleistocene (details in Table 1).

**Table 1.**
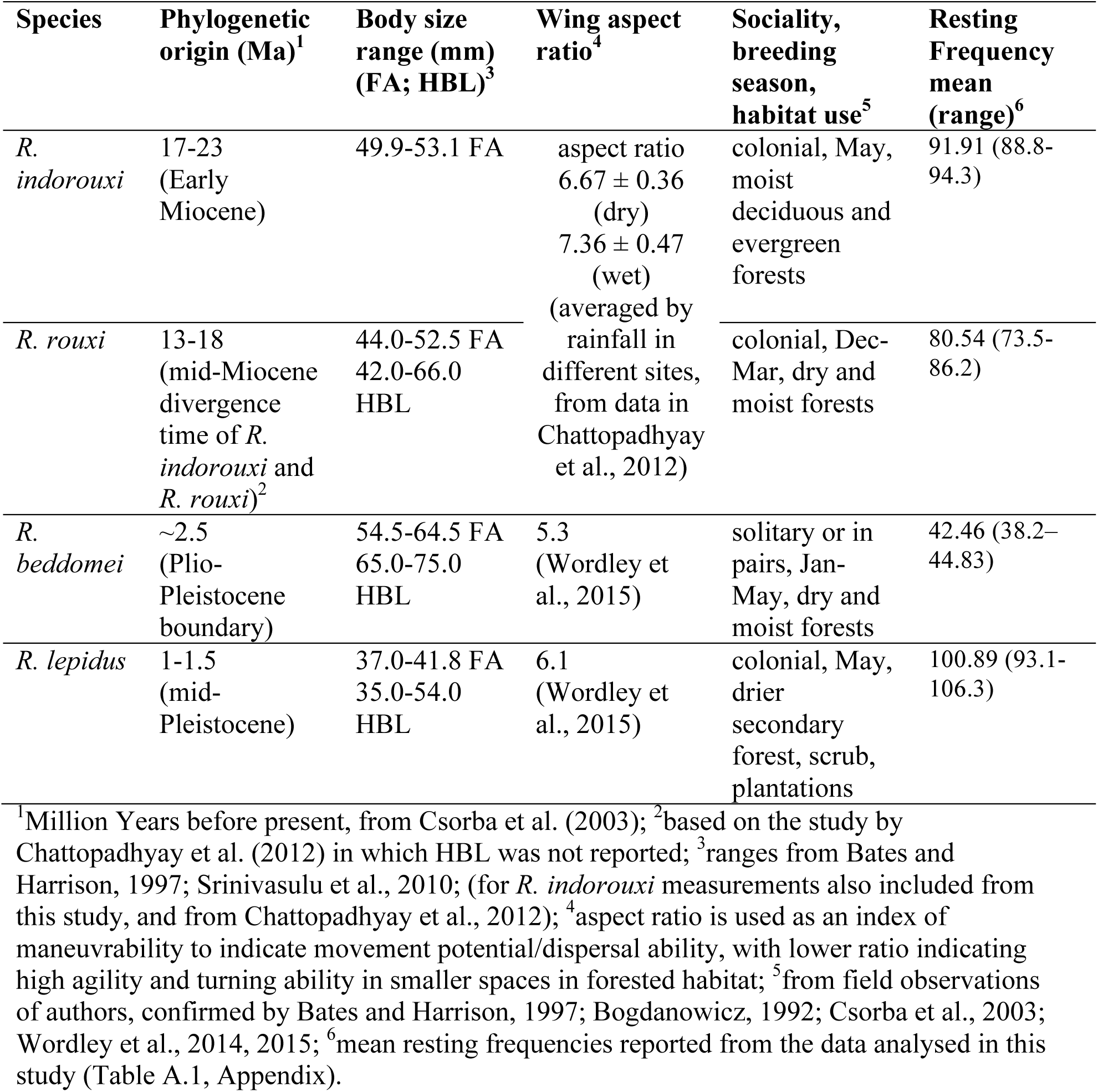
A summary of *Rhinolophus* species studied, with their phylogenetic information, morphometric measurements, life-history traits, and resting frequencies.

### Sampling and analyses of Resting Frequencies of *Rhinolophus* species

Bat species were identified with standard morphological keys (e.g. Bates and Harrison, 1997). We analysed recordings made either from captured handheld bats, or individual bats focally sampled in their roosts – together referred to as “stationary” bats throughout the text. The latitude and longitude of roost sites were logged in a Global Positioning System (Garmin eTrex vista H). From their roosts, bats were captured with a hand-net. They were allowed to rest and stabilize post-capture, recordings were made to obtain data on resting frequencies (RF), and bats released at their roosts. When capturing bats was not possible, we conducted recordings of individual adult-sized bats roosting solitarily. Distances from 0.3 to 1 m were maintained between stationary bats and the detector to record calls at optimal sound source levels. This helped avoid artefacts from reflections or frequency changes from acoustic compensation, for accurate assessment of RFs (Jacobs et al., 2017; Jiang et al., 2010b, Schuchmann and Siemers, 2010). We recorded 7820 resting calls, from 2298 sequences, over 290 h, for the four *Rhinolophus* species, from 24 roosting sites in 5 localities (Table A.1, Appendix). An example of *Rhinolophus* calls is presented in Fig. 2. We used only RF calls for final analyses, to avoid the influence of Doppler shift compensation (e.g. Jiang et al., 2015).

**Figure 2.**
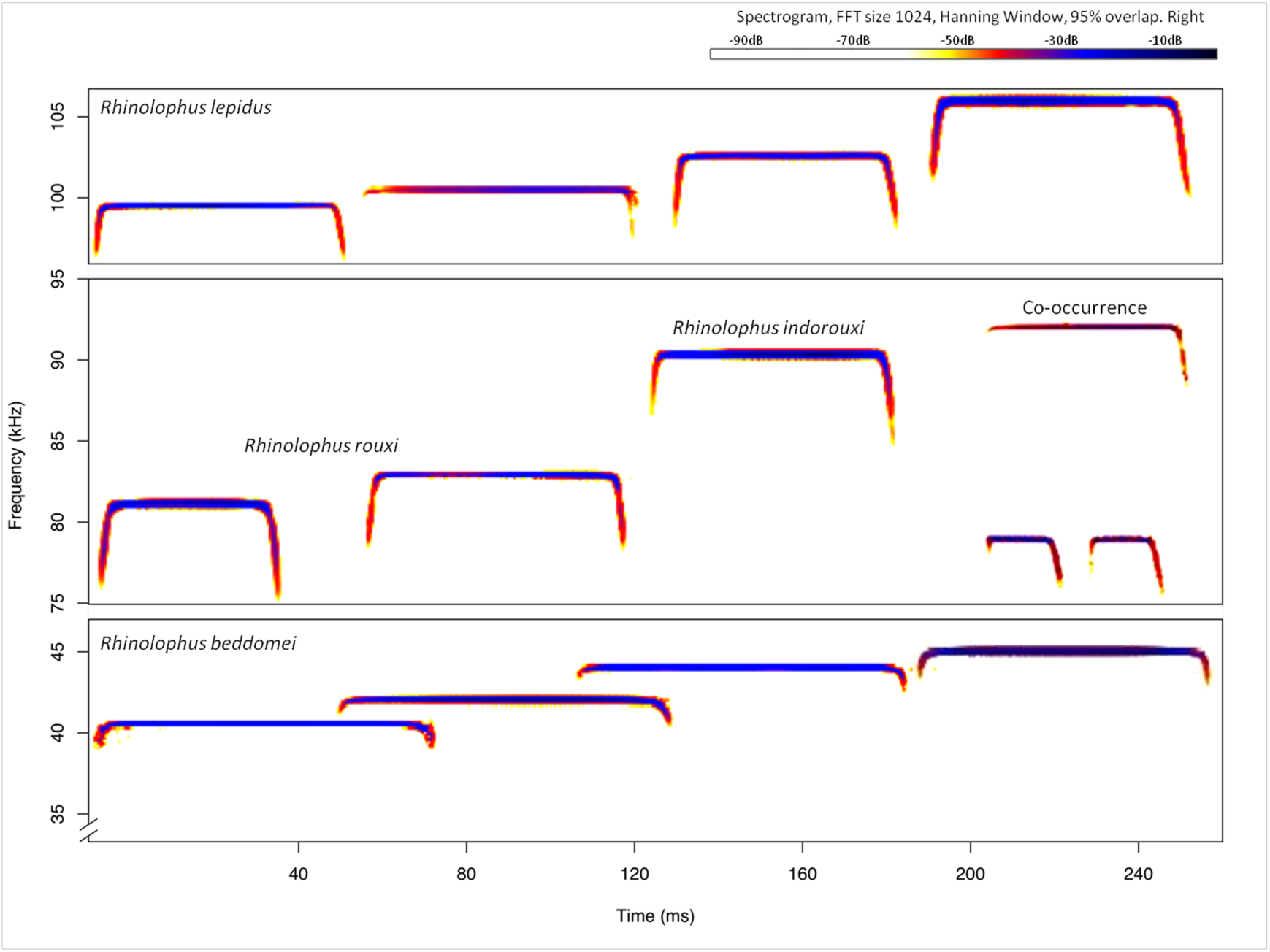
Representative echolocation calls of the four *Rhinolophus* species. Top and bottom panels show diversity of acoustic frequencies in *R. lepidus* and *R. beddomei* respectively. Middle panel shows the diversity for *R. rouxi* and *R. indorouxi* when they occurred singly and co-occurred (left to right).

All recordings were made with an ultrasonic bat detector (D240X, Pettersson Elektronik AB, Uppsala, Sweden) set at high-gain automatic detection settings with 10× time expansion, and recordings were stored in an external recorder (Edirol R-09HR; Roland; www.edirol.net). The sampling rate of the detector was 307 kHz with detector range from 10 to 120 kHz. This sampling rate was adequate in resolution for accurate estimation of RF characteristics of the species we recorded (RFs ranging between 38 and 106 kHz). Given our financial constraints and for comparability with other studies (e.g. Wordley et al., 2014, 2015), this was considered the best available detector. We compared calls in .wav format with calls in .mp3 format. As we did not find any difference in call parameters estimated (resting frequency, pulse duration, amplitude), for this work we recorded calls in .mp3 format to reduce file size for storage (Brigham et al., 2004), which was necessary in remote field conditions. Recorded .mp3 data files were converted to .wav format in GOLDWAVEsoftware (www.goldwave.com) for analyses in the software BATSOUNDPROv.3.32 (Pettersson Elektronik AB, Uppsala, Sweden, www.batsound.com). Recordings were analyzed using sampling frequency of 44.1 kHz at FFT size of 1024 samples at 95% overlap in a hanning window (Xu et al., 2008). We measured the Resting Frequency (RF) as the CF-component of FM-CF-FM calls of stationary *Rhinolophus* bats from the power spectrum in sonograms (Stoffberg et al., 2012). Call duration and inter-pulse intervals (IPI) of calls in milliseconds (ms) were also measured to aid in call selection of individual roosting bats for analysis (Brigham et al., 2004; Xu et al., 2008). Avoiding the first ten recorded pulses of stationary bats (Jacobs and Bastian, 2018) from sequences with unambiguous individual calls having high signal-to-noise ratio, we extracted mean RF (the dominant 2^nd^ harmonic), minimum RF, and maximum RF, for each individual roost site. Numbers of representative RFs for statistical analyses were as follows: *R. indorouxi* (n=16), *R. rouxi* (n=28), *R. beddomei* (n=10), *R. lepidus* (n=12).

### RF data from literature review

We compiled published data on RFs of the four *Rhinolophus* species. Again, we excluded any information on *R. pusillus* owing to limited data and possible acoustic misidentification with *R. lepidus* (Appendix A.). We used strict criteria in selecting studies to augment our RF data, which were: a) reported RFs were from hand-held bats only, b) detectors used for the studies had sampling frequencies comparable to our detector, c) recording and analysis methods were consistent in selection of calls, and d) species identification was consistent. We excluded any data from unverifiable sources. We also verified field records with laboratory recordings reported by other studies (Appendix A). Data on resting frequencies were compiled from published journal articles (n=19), reports (n=2), doctoral and masters’ dissertations (n=6), and website reports, from a total of 16 localities in the WGSL. A list of data sources and numbers of bats recorded is provided in Table A.1 (Appendix). In spite of these checks, we are aware that “measurement errors” would exist, and cannot be estimated for the different devices and methods used (Adams et al., 2012; Kelly, 2008; Goerlitz, 2018). As they would affect our ability to compare multi-source RF data in the same statistical analysis, we used the data only to qualitatively assess patterns that could hint at acoustic divergence.

We extracted minimum, maximum, and mean of RFs for analysis from reported summary statistics of resting frequencies across these locations, resulting in the following representative RF samples for analysis: *R. indorouxi* (n=12); *R. rouxi* (n=19); *R. beddomei* (n=14); *R. lepidus* (n=10). From our own data and published data, we obtained 121 RFs for statistical analyses. Our sample selection led to substantial data reduction, but allowed for qualitative analysis of the entire range of variability of RF signatures for individual roosts. It also helped avoid pseudo-replication of repeated calls sampled from the same sequence. This led to conservative yet consistent assessment of environmental effects on RF.

### Environmental and bioclimatic variable selection

We extracted bioclimatic data for all locations from where average resting frequencies were obtained. Data on RH, i.e. Relative Humidity (www.en.openei.org) were obtained at 1° x 1° resolution at 10 m above the earth’s surface (NASA SSE 6.0 2007). We downloaded precipitation and temperature data from BIOCLIM (http://www.worldclim.org/bioclim), and elevation and terrain data from the Shuttle Radar Topography Mission (www.cgiar.org). We also extracted NDVI (Normalized Difference Vegetation Index for vegetation greenness) from MODIS satellite images (250 m × 250 m resolution) (www.earthexplorer.usgs.gov).

From these data we characterized monsoonal duration as highly pulsed or relatively uniform. All variables were averaged over a 1^°^ × 1^°^ resolution to match the humidity data. Changes in humidity along latitudinal and longitudinal gradients, corresponding to the spatial scale of the pre-defined biogeographic-climatic zones, were estimated (e.g. Krishna Prasad et al., 2008). We used Mantel tests (Goslee and Urban, 2007) to detect correlations between pair-wise differences in relative humidity with differences in latitude and longitude. The strength and statistical significance of the Mantel test coefficients (Mantel’s r>0.25, p<=0.001) were the criteria for choosing whether latitude and longitude should be added as spatial covariates to our regression models. To detect correlations between humidity and other environmental variables (elevation, rainfall, NDVI, temperature, latitude, longitude; Table 2), we used Spearman’s rank correlation tests. We did not expect spatial effects on correlation strength due to the coarse resolution of data used. Elevation was not correlated significantly with humidity. We chose relative humidity (RH) and elevation (ELEV) as the two main explanatory variables, and rainfall (RAIN) as an additional covariate in regression models.

**Table 2.** Variable selection based on Spearman’s rank correlation tests, showing the strength and statistical significance of correlations between Relative Humidity (RH) and other environmental variables, along with latitude and longitude, for all species’ locations. The table lists variables that were strongly and consistently correlated (p<0.05) with RH and therefore not used in regression models. RH consistently decreased northwards and increased eastwards, and was strongly correlated with topography and NDVI. Only elevation, rainfall, and RH were used in regression models.

| Environmental variables | Correlation with Relative Humidity (species-wise) |  |  |  |
| --- | --- | --- | --- | --- |
|  | *Rhinolophus indorouxii* | *Rhinolophus rouxii* | *Rhinolophus beddomei* | *Rhinolophus lepidus* |
| Topography | 0.35 ($p=0.001$) | -0.52 ($p<0.001$) | -0.49 ($p<0.001$) | -0.375 ($p<0.001$) |
| NDVI | 0.24 ($p=0.03$) | 0.45 ($p<0.001$) | 0.89 ($p<0.001$) | 0.42 ($p<0.001$) |
| Latitude | -0.86 ($p<0.001$) | -0.96 ($p<0.001$) | -0.85 ($p<0.001$) | -0.96 ($p<0.001$) |
| Longitude | 0.73 ($p<0.001$) | 0.82 ($p<0.001$) | 0.32 ($p<0.001$) | 0.92 ($p<0.001$) |

### Species-wise information on body size and humidity

From the reviewed literature and our own measurements, we extracted data on RF, body size, and humidity at the approximate geographic coordinates and habitat description of each recorded location. We obtained body size data (forearm length in mm) for 30 individuals across the four bat species. Locations from which we obtained body size data were grouped into dry and humid habitats, based on bioclimatic variables. We compared body size data for each species across these habitats to qualitatively test the predictions of the James’ rule for all species.

### Observations on co-occurrence of *R. rouxi* and *R. indorouxi*

We identified roosts where these sibling species co-occurred. We assessed differences in the RFs of these species in roosts where they occurred together, and in single-species roosts. We then assessed their distribution and foraging habitat use (based on night-time acoustic recordings) in relation to elevation, humidity, and habitat types used.

## Statistical analyses

### Prediction of resting frequency based on relative humidity and elevation

We used exploratory and graphical analyses to assess variation in resting frequency with respect to annual relative humidity, elevation, and annual rainfall at bat roosts and other recording locations. For each species, we ran separate sets of candidate models using Generalized Least Squares (GLS) regression models (Dormann et al., 2007) fitted with a restricted maximum likelihood procedure to model spatially structured variation in frequency, humidity, elevation, and rainfall. All predictor variables were scaled in the models we tested. To detect the nature of spatial correlation we empirically estimated the sill and range parameters of semivariograms. Spatial correlation based on latitude and longitude (projected to the Universal Transverse Mercator system, zone 43-N for estimating Euclidean distances) was estimated from an autoregressive smoothing parameter (moving average parameter) or parameters of an exponential model (corAR1 and corExp families, respectively). We compared the log-likelihood (measure of fit) of GLS models with ordinary least square (OLS) models that did not account for spatial structure in environmental variables (Dormann et al., 2007). The Akaike Information Criterion (AIC) was used to select the most parsimonious and best fitting of all candidate models (based on Burnham and Anderson, 2002). All analyses were conducted in the packages ‘ecodist’ (Goslee and Urban, 2007) and ‘nlme’ in R 3.2.3 (R Core Team, 2015).

The data used to fit the models had unknown measurement errors. The RF data (response variable) for each bat species were compiled from our own recordings and published data sources with comparable recording devices, however, the exact specifications of all devices used in the published papers were not available to us. Due to this issue, it was not possible to assess the contribution of relative measurement errors (across devices) to our inferences about RF responses. Therefore, we interpreted only the sign, and not the magnitude of RF responses (slope parameter estimates for the GLS models). Our reasonable assumption was that, because all recording devices had comparable sampling rates, any measurement errors would not hinder our ability to detect the tendency of RF responses, but only affect the numerical estimation of strength of response.

### Variation in resting frequency across biogeographic zones

We systematically examined RF variations of each species within and across the seven biogeographic zones (study area), using exploratory graphical analyses in the R ‘base’ package. We also compared RFs across two major biogeographic barriers: 1) north and south of the Palghat Gap within India’s Western Ghats, and 2) in India and Sri Lanka, with Wilcoxon’s rank-sum tests.

### Relationships between body size, frequency, and humidity

We checked if the correlations between body size (forearm length), resting frequency, and humidity were statistically significant, with Spearman’s rank correlation tests (due to its statistical power despite small sample sizes) for each species. Due to limited data, we could only qualitatively assess whether species’ body sizes followed predictions of the James’ rule.

## Results

### Geographic gradients in relative humidity

Humidity decreased with increasing latitudes along the Western Ghats (Mantel’s r = 0.81, p=0.001). Higher and more uniform humidity profiles characterized the western slopes of the Western Ghats escarpment southward from 15° latitude, at elevations around 600-900 m a.s.l. With longitude, humidity consistently reduced eastward as rainfall reduced (Mantel’s r = 0.29, p=0.003), except over the Nilgiri, Annamalai, and Palni high-elevation massifs.

### Spatially corrected effects of atmospheric relative humidity on resting frequency

Effects of Relative Humidity (RH), elevation, and rainfall varied considerably as predictors of Resting Frequency (RF) of the four species, as per the spatial regression models. For *R. indorouxi*, elevation and RH both predicted RF almost equally well; for *R. rouxi*, humidity was the main predictor of RF. For *R. beddomei*, elevation, rainfall, and humidity were all important predictors (Table 3, Fig. 3). But for *R. lepidus*, rainfall was the best predictor of RF. Our hypothesis that RF will be lower at higher RH was supported only by *R. indorouxi* and *R. rouxi* (negative slopes). Positive relationships with RH were found for *R. beddomei* and *R. lepidus* (Table 3; Fig. 3). Effects of elevation on RF were negative for *R. beddomei* and *R. rouxi*, but positive for *R. indorouxi*, a species occurring at higher altitudes than its sibling *R. rouxi* (Table 3; Fig. 3). Indirectly, elevation and rainfall could have influenced humidity effects on these species.

**Table 3.**
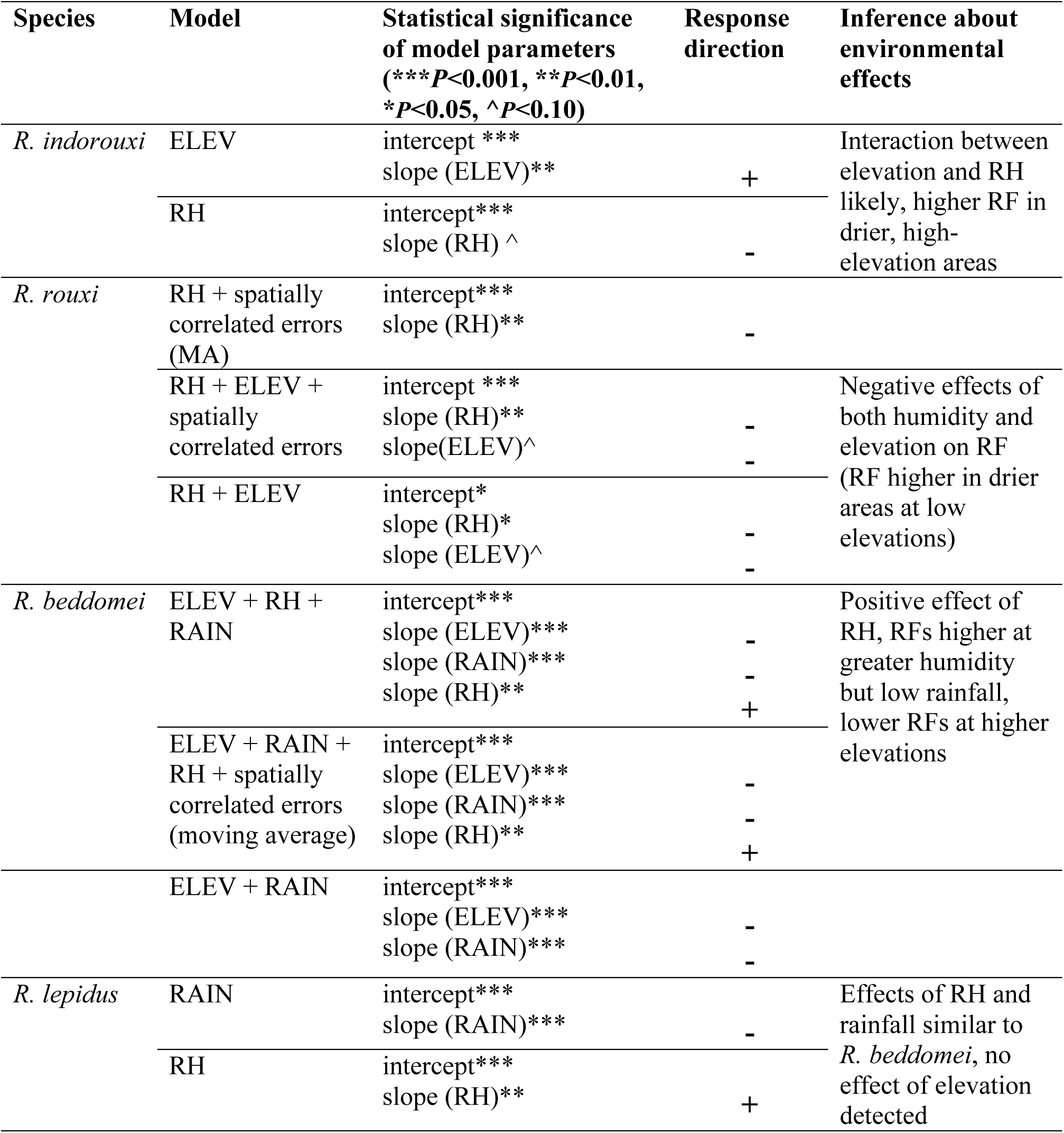
Model selection results and estimated effect sizes of relative humidity (RH), elevation (ELEV), annual rainfall (RAIN), and spatial effects for the *Rhinolophus* bats studied (MA coefficient = Moving Average smoothing parameter in Generalized Least Squares models). Model selection was based on log-likelihood and residual standard errors, as well as δ-AIC comparisons. Signs of responses (response direction) of RF and levels of statistical significance (p-values) are shown for the best models’ outputs for each species.

**Figure 3.**
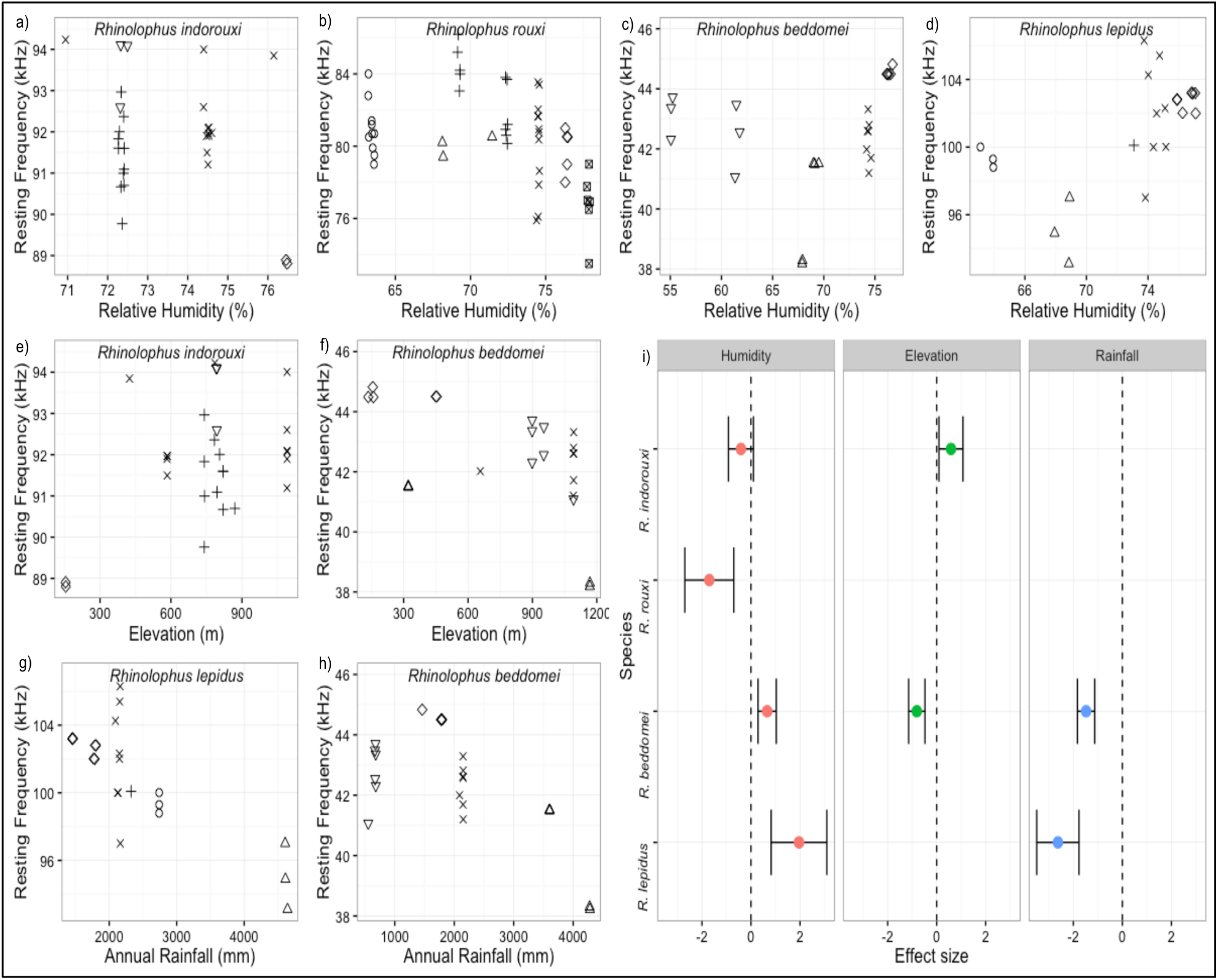
Humidity had contrasting correlations with RF: (a-b) for *R. indorouxi* and *R. rouxi*, the hypothesis that RF will reduce with increasing humidity was supported, but for (c-d) *R. beddomei* and *R. lepidus*, a different pattern was noted. *R. indorouxi* and *R. beddomei* RFs were higher and lower respectively, at higher elevations (e-f), and *R. lepidus* and *R. beddomei* RFs were positively correlated with annual rainfall (g-h). A comparison of effect sizes of humidity, elevation, and rainfall on the resting frequencies (RF) of four *Rhinolophus* species based on Generalized Least Square (GLS) models are provided in (i). Plot symbols indicate biogeographic zones along latitudes of the Western Ghats-Sri Lanka region (see Fig.1 for abbreviations): open circles (N.G), triangles (G-CV), plus signs (CV-PG), multiplier signs (PG-SG), diamonds (S.SG), inverted triangles (P.EG), and squares with diagonals (SL).

### Variation in resting frequency across biogeographic zones

RFs of the four *Rhinolophus* species across the seven biogeographic zones showed no consistent differences (Fig. 4). For *R. beddomei* and *R. lepidus*, RFs were significantly lower than the overall mean in the high-rainfall region of the central Western Ghats (Goa-Cauvery valley region) and higher in lower rainfall areas south of the Shenkottah Gap (Fig. 4). RF of the Sri Lankan subspecies *R. rouxi rubidus* was significantly lower than *R. rouxi* recorded in the Western Ghats (Wilcoxon’s rank-sum test; W=23, p=0.0003). RF did not vary much across different zones for *R. rouxi* barring a slightly higher median RF in the zone north of the Palghat Gap, where it co-occurred with *R. indorouxi. R. indorouxi* RFs were lower than the overall mean only in the zone south of the Shenkottah Gap, where *R. rouxi* also occurred close to *R. indorouxi*. These results suggested that environmental variables might have had a greater effect on RF than biogeographic barriers (Fig. 4).

**Figure 4.**
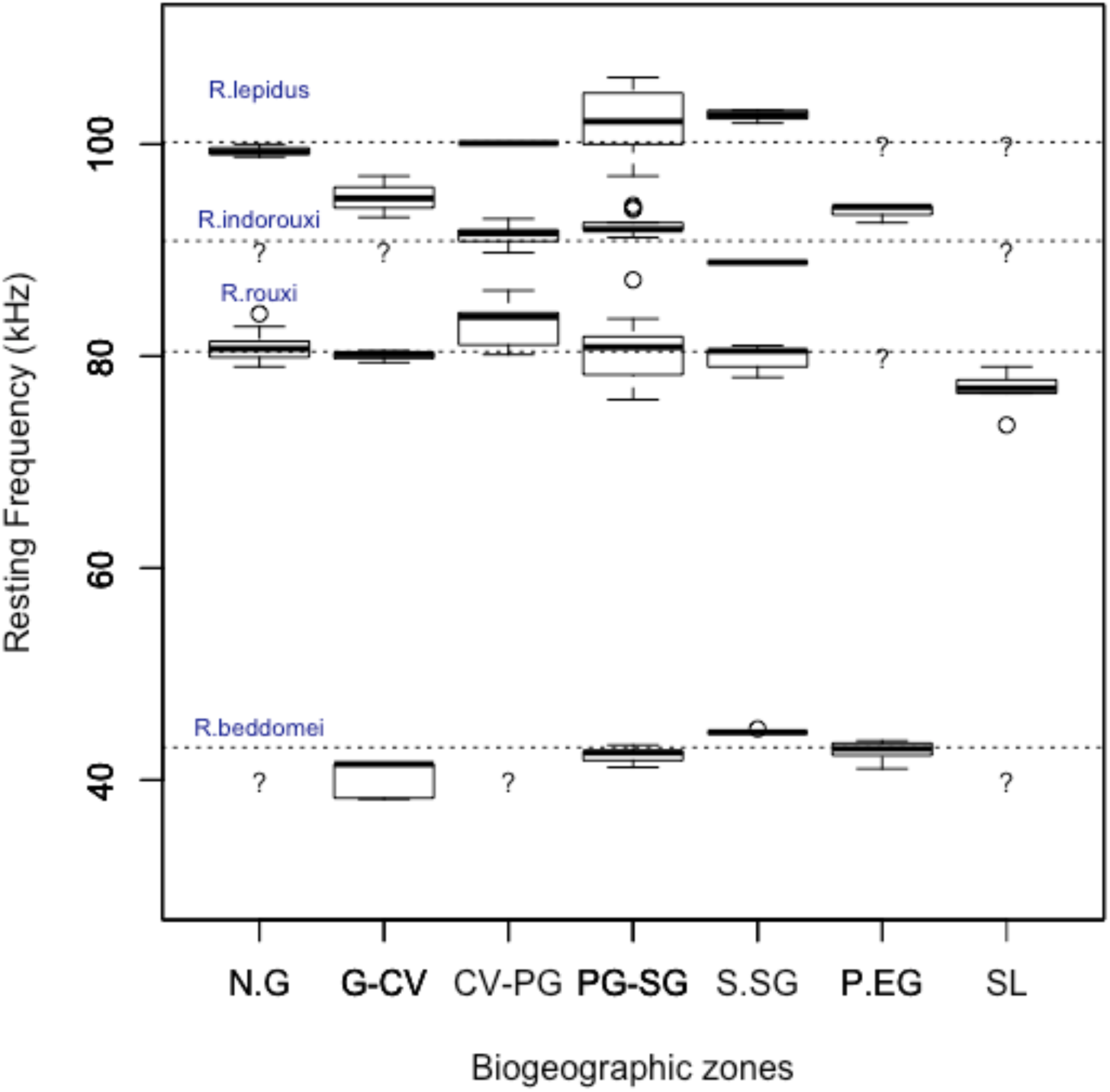
Variation in resting frequencies of four *Rhinolophus* species across biogeographic zones of the Western Ghats-Sri Lanka region (see Fig. 1 for zone abbreviations). Question marks indicate zones with missing data; *R. lepidus* is considered absent from Sri Lanka (see Methods section).

### Body size, humidity, and resting frequency

All species did not uniformly follow the James’ rule. Forearm lengths of individuals in hot and humid areas were lower than of bats in cool and dry areas for *R. indorouxi* and *R. lepidus*, but an opposite trend was noted in *R. beddomei*. In general, *R. indorouxi* (the larger sibling species) occurred at cool and dry high-elevation sites in the WGSL and peninsular India, and the smaller *R. rouxi* occurred in the warmer lowlands (Fig. 5). Individuals of the Sri Lankan (insular) subspecies/populations of *R. rouxi* and *R. beddomei* were smaller than those from hot and humid regions of peninsular India. But the correlation between body size (forearm length) and humidity was not statistically significant for any of the species, at α=0.05. Forearm length and RF were also not correlated. The only exception was *R. indorouxi*, where larger individuals had higher RFs, indicating an effect of humidity perhaps not influenced by body size (Table 4). For other species, signs of the correlation coefficients were negative, pointing to an interaction of humidity and allometric effects on RF.

**Table 4.** Correlations (Spearman’s rho) of humidity and resting frequency with forearm length (body size) in four species of *Rhinolophus* across the Western Ghats-Sri Lanka biodiversity hotspot. No significant correlations were found, except a positive correlation of forearm length with frequency in *R. indorouxi* (\**P*<0.05).

| Species | n | Forearm length ~ Humidity | Forearm length ~ Resting Frequency |
| --- | --- | --- | --- |
| *Rhinolophus indorouxi* | 10 | rho = -0.29 (p=0.42) | rho = 0.68 (p=0.03)* |
| *Rhinolophus rouxi* | 8 | rho = 0.42 (p=0.35) | rho = -0.64 (p=0.18) |
| *Rhinolophus beddomei* | 6 | rho = 0.22 (p=0.55) | rho = -0.14 (p=0.78) |
| *Rhinolophus lepidus* | 6 | rho = -0.67 (p=0.14) | rho = -0.70 (p=0.23) |

**Figure 5.**
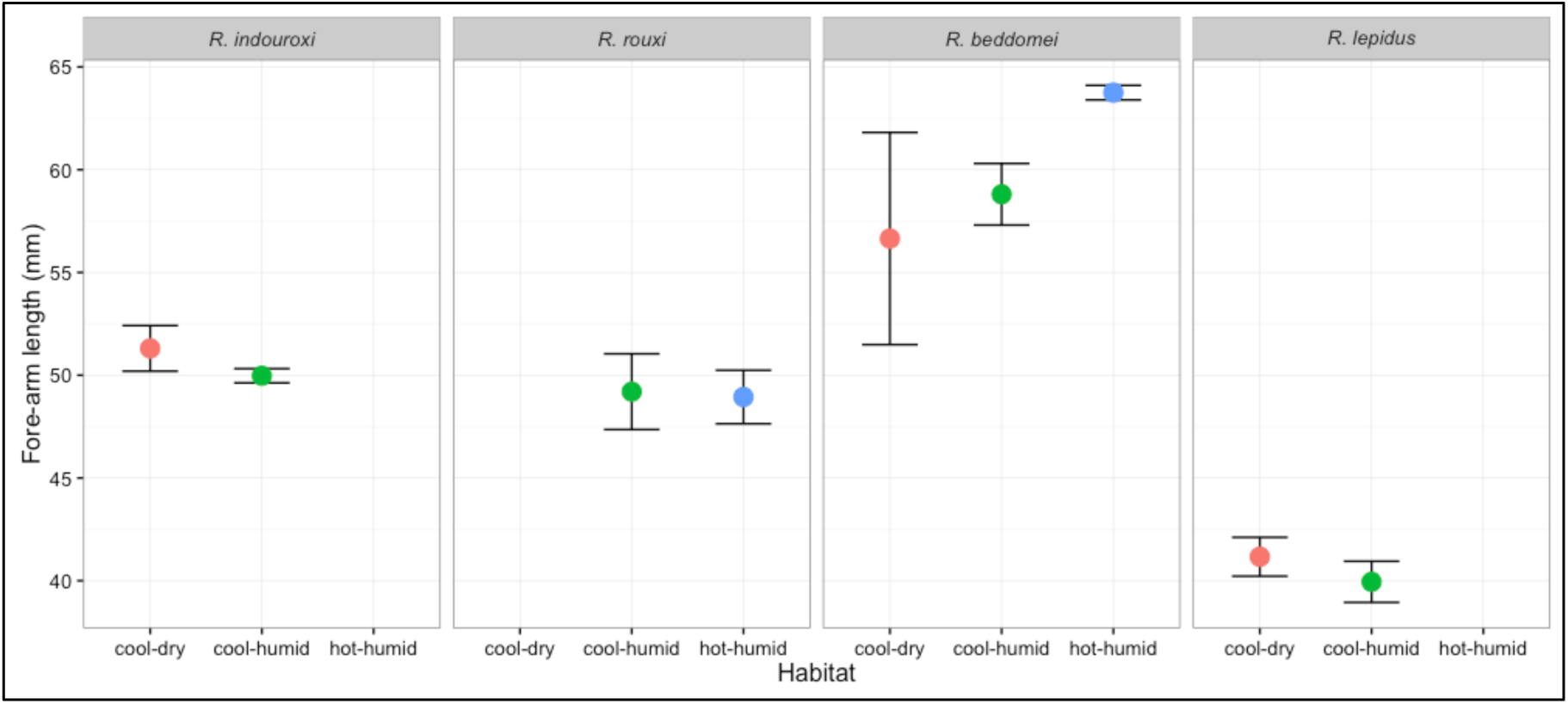
Preliminary comparisons of forearm lengths across cool-dry, cool-humid, and hot-humid habitats, to test the James’ rule for the four *Rhinolophus* species. *R. indorouxi* and *R. lepidus* showed larger forearm lengths in cool, dry habitats as compared to humid habitats. *R. beddomei* showed an opposite pattern.

### Acoustic variation and co-occurrence of sibling species of the *R. rouxi* complex

*R. indorouxi* and *R. rouxi* distribution overlapped along the 600-900 m elevation band along the western escarpment of the Kerala-Tamil Nadu water divide, along the northern edges of the Nilgiri and Anamalai high-elevation massifs. In shared roost sites (at Wayanad and Parambikulam), RFs of the sibling species were strongly non-overlapping (maximum RF=84 kHz for *R. rouxi* and minimum RF=92 kHz for R. *indorouxi*) as compared to single-species roosts, where the range of *R. indorouxi* RF was 89-94 kHz, and that of *R. rouxi* was from 73-85 kHz. In shared roosts, the species roosted in two separate groups. Clear spatial segregation was also observed in foraging habitats near these roosts. At these sites, we recorded *R. indorouxi* only in cool, moist foraging habitats such as evergreen forests or shade coffee, and *R. rouxi* only from hotter, drier habitats (deciduous forests, teak plantations). These species foraged in wet and dry forests separated only by 15-20 km in the Shenduruney landscape, in the Agasthyamalai mountain ranges. In mid-elevation ‘co-occurrence zones’, calls of the two species could thus be affected by local humidity.

## Discussion

Our preliminary study identified potentially significant, but variable, influences of relative humidity and elevation on acoustic divergence in the four *Rhinolophus* species studied in the WGSL region. We also did not find any consistent variation in species’ RFs across biogeographic zones. Not finding any consistent differences across biogeographic zones is a significant result. It suggests stronger effects of environmental variability on acoustic divergence in WGSL Rhinolophids owing to adaptive processes (sensory drive) than due to gene flow barriers (Wilkins et al. 2013, Jiang et al. 2015). Indeed, the effects of humidity were consistent locally and regionally, influenced by rainfall and elevation to variable extents. Qualitatively, *R. indorouxi* and *R. lepidus* obeyed predictions of the James’ rule, whereas *R. beddomei* showed an opposite pattern. This suggested differential effects of humidity and body size on RF, but correlations between them were not statistically significant (mostly due to limited sample size). More than body size, changes in specialized organs, such as the nose-leaf, nasal capsule, or pinna size might be related to RF variation (Armstrong and Coles, 2007; Wu et al., 2015). Differences in these structures along environmental and biogeographic gradients should be measured in future studies to confirm our preliminary findings.

Our results broadly resemble findings from *Rhinolophus* bats in Africa (Jacobs et al., 2017; Jacobs and Bastian, 2018; Mulaleke et al., 2017; Mutumi et al., 2016; Stoffberg et al., 2012), on correlated environmental and phylogenetic factors influencing divergence (Jones, 1997; Sun et al., 2013). The heterogeneous effects of humidity on the four species’ RFs merit discussion. We propose that RF responses to humidity might differ because of evolutionary history, i.e. in relation to the palaeo-climatic conditions in which the four *Rhinolophus* species diverged. Chattopadhyay et al. (2012) attributed the divergence of *R. indorouxi* and *R. rouxi* to Miocene drying in southern India. Moreover, these sibling species evolved from a basal lineage that probably originated in the peninsular Indian craton (Csorba et al., 2003), and perhaps had longer time unto speciation through acoustic divergence and potential reproductive isolation (e.g. Kingston et al., 2001; Mao et al., 2010). This was unlike *R. beddomei* and *R. lepidus*, which dispersed into India from the Malayan region and Indochina respectively, and diverged from their ancestors more recently, in the Pleistocene. *R. beddomei* evolved at the time of the Early Pleistocene (2.5 Ma) when monsoonal climate intensified (Csorba et al., 2003; Kale et al., 2003; Zhang et al., 2013). *R. beddomei* prefer evergreen forests (Bates and Harrison, 1997; Wordley et al., 2015), which might explain their positive correlation of RF with humidity. *R. lepidus* colonized peninsular India about 1.5 Ma (Quaternary period), when aridification effects were strong and rainfall was pulsed (Kale et al., 2003; Gower et al., 2016). Rhinolophids of the *R. pusillus* sub-group (to which *R. lepidus* belongs), underwent major range expansions in this period (Dejtaradol, 2009; Jiang et al., 2010b). Thus, the positive correlation of RF and humidity for *R. lepidus*, which is a bat of semi-arid biotopes and deciduous scrub forests, is likely a sampling artefact, given limited data from the WGSL. For *R. lepidus* and *R. beddomei*, the contrasting effects of RH and rainfall (Figure 3) might be related to the seasonally concentrated rainfall distribution and temperature differences in the central Western Ghats. *R. lepidus* frequencies were distinctly lower in drier regions of the central Western Ghats, suggesting acoustic divergence between dry and wet forest populations of the WGSL. The negative correlation of its RF to rainfall supported this. Possible reasons proposed for the positive response of *R. pusillus* to humidity by Jiang et al. (2010b) could apply to *R. lepidus*, where bats might increase RF to overcome effects of rain noise for better signal transmission, or to capture tiny prey such as mosquitoes or flies (Csorba et al., 2003; Dejtaradol, 2009).

For the sibling species *R. indorouxi* and *R. rouxi*, we found resting frequencies forming a “phonic continuum” covering the frequency range from 73 to 94 kHz across the WGSL. The *R. rouxi* complex might thus have more cryptic species than currently known (Csorba et al., 2003; Thomas, 1997). *R. indorouxi* and *R. rouxi* co-occurred along a distinct “zone of contact” (600-900 m a.s.l. along the Western Ghats escarpment). This could indicate either a ‘hybrid zone’ formed by secondary contact of the two species (Mao et al., 2014; Sun et al., 2016). It will be interesting to study the degree of introgression and hybridization between the sibling species along these zones of contact. As *R. rouxi* likely diverged from *R. indorouxi*, a parapatric mode of speciation is also likely (Fitzpatrick et al., 2009; Mallet et al., 2009; Rundle and Nosil, 2005). This zone overlaps a part of the elevation range in which parapatric speciation in bush-frog lineages was recently described (Vijayakumar et al., 2016). Interestingly, co-occurrence of these sibling species was recorded only in three wet-dry transition zones that have recorded stable rainfall over the last four decades (Rajendran et al., 2012). These sites were along the ridge of the southern Western Ghats (Wayanad, Parambikulam, Shenduruney), and would be influenced by orographic effects on rainfall caused by adjacent high-elevation massifs. *R. indorouxi*, when not co-occurring with *R. rouxi*, was found mostly above this zone in cooler high-elevation areas (900-2000 m). Below this zone (150-600 m) it was rare and restricted to humid and moist forests. In contrast, *R. rouxi* occurred mostly in dry forests from sea level up to 1000 m, above which it was rarely encountered, except in Sri Lanka (where *R. indorouxi* is absent). *R. indorouxi* is the larger species of the two and also has a larger RF. Food resource partitioning and prey availability in different types could allow for such deviation of RFs from allometric predictions (Kelly, 2008, Shi et al., 2009; Lee et al., 2012). RFs of the insular subspecies *R. rouxi rubidus* were lower than Western Ghats *R. rouxi rouxi*, in spite of no clear differences in forearm lengths. This suggests that Sri Lankan *R. rouxi rubidus* RFs were different due to character displacement or environmental effects (Russo et al., 2007). A similar comparison remains wanting for *R. beddomei sobrinus* of Sri Lanka, which are much smaller than Indian specimens (Csorba et al., 2003).

Figure 6 conceptually summarizes our findings on a general pattern of co-occurrence of the sibling species of the *R. rouxi* complex. Interestingly, we observed that *R. indorouxi* and *R. rouxi*, even when using the same roost, clustered in non-overlapping groups (based on detected frequency variations within and between groups), suggesting assortative mating (Park et al., 1996). As per the general prediction by Russo et al. (2007), we saw that RFs showed strong separation in roosts where these sibling species co-occurred. Separate foraging sites used by these species was another observation matching what Salsamendi et al. (2012) reported as the reason for coexistence of sympatric sibling *Rhinolophus* species. Together, these observations provide an environmental and geographic context to the genetic findings of Chattopadhyay et al. (2012).

**Figure 6.**
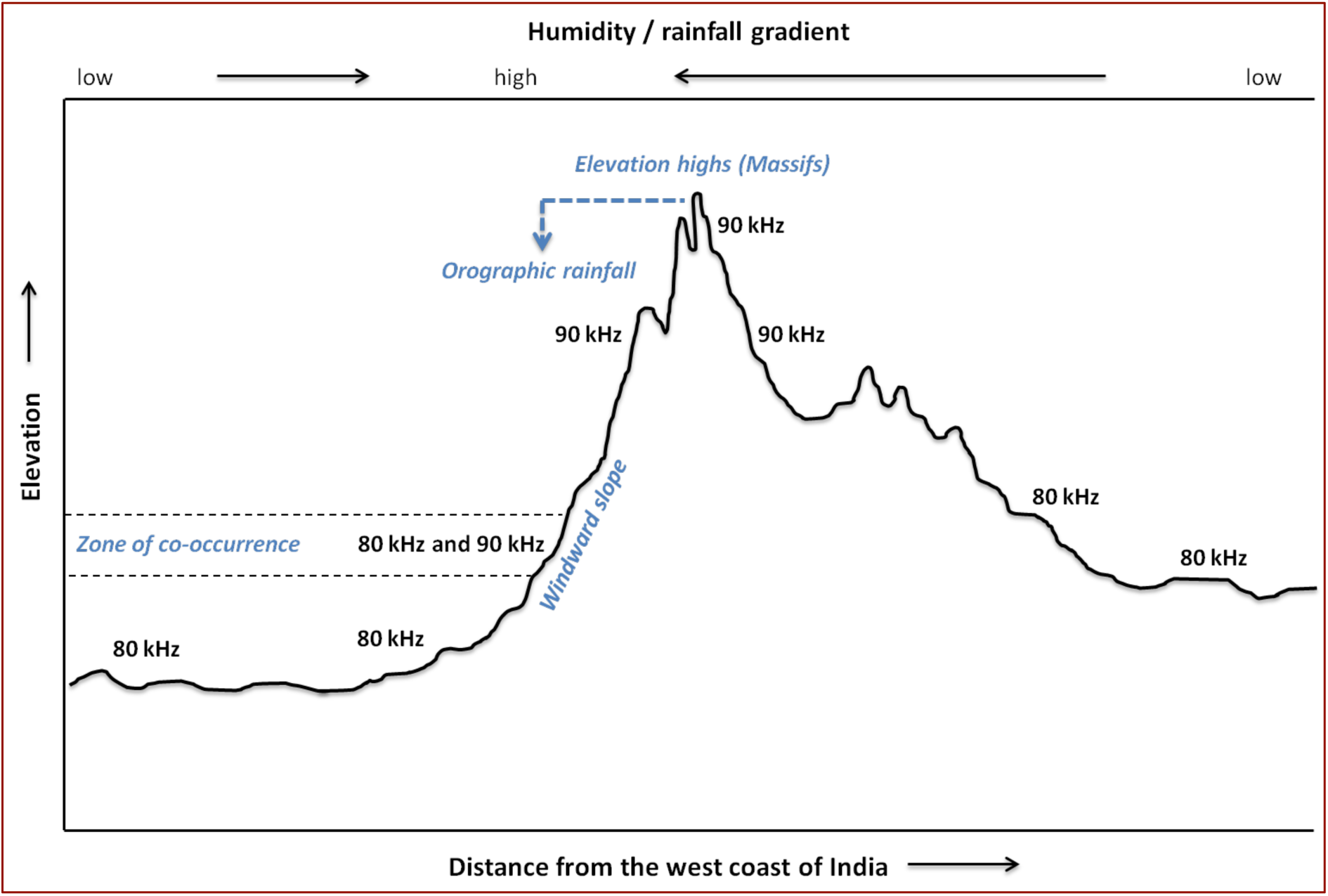
Schematic of the distribution of the sibling species *R. indorouxi* (RF 92±3 kHz) and *R. rouxi* (RF 80±6 kHz) along a transect drawn from the coastline of west India through the Western Ghats escarpment, elevation highs, and plateau regions of southeastern peninsular India. Zones of co-occurrence appear to exist along western windward slopes of the WG, in areas with high rainfall and humidity from orographic effects. At lower elevations to the west and east of this zone, only *R. rouxi* was detected, and at high elevations only *R. indorouxi* was detected. We predict that this pattern occurs in other areas of the WG with similar relief and humidity profiles. [This diagram for representational purposes only.]

Our results emphasize the importance of environmental variables in driving cryptic speciation of *Rhinolophus* that needs to be tested with future work on gene flow (Jiang et al., 2015; Kingston et al., 2001; Russo et al., 2007). The study thus contributes some insights to potential mechanisms of environmentally mediated acoustic divergence. Our ideas should be tested with combined phylogenetic and acoustic studies, as more consistent and comparable data on these species become available in the future.

*Rhinolophus* bats are sensitive to deforestation, fragmentation, and microclimatic changes (Struebig et al., 2008; Wordley et al., 2015). Anthropogenic barriers to gene flow through land-use and climate change could further drive acoustic variation and co-occurrence patterns (Jacobs et al., 2017; Purushotham and Robin, 2016; Sedlock and Weyandt, 2009; Xu et al., 2008). Studies on genetic/cultural drift or sexual selection should interpret acoustic differences in *Rhinolophus* bats by accounting for environmental change impacts in rapidly changing biotopes like the WGSL.

## Acknowledgements

We are indebted to Mahesh Sankaran, National Centre for Biological Sciences, Bangalore, for his constant support and providing on loan the acoustic detectors used in our study. We are grateful to Lasse Jakobsen and Peter Madsen for their critical advice in bat acoustic theory and practical analysis methods. We thank the officials and staff of the Kerala and Karnataka Forest Departments for providing research permits and logistical support. Assistance in fieldwork by Shinu Jacob, Arshad, Gaurav Kalyani, Ashwin Varudkar, Kaustubh Deshpande, and others, is sincerely acknowledged. Thanks are due to all authors whose published and reported work helped us conduct our preliminary analysis of acoustic divergence at a large scale. This work was partly funded by the Rufford Foundation Small Grant (ref: 9686-1), United Kingdom, and a small grant from the World Wildlife Fund (WWF), India, to Kadambari Deshpande.

## Appendix

**Table A.1.** Data sources and details of locations from which *Rhinolophus* resting frequency data were collected and analyzed.

| Biogeographic/<br>climatic<br>zone | Location<br>(no. of roosts surveyed) | Forest<br>type | Bat species<br>recorded | Sample<br>size<br>(no. of<br>focal bats) | Data sources |
| --- | --- | --- | --- | --- | --- |
| Primary data on RFs collected by authors |  |  |  |  |  |
| N.G | Barapede (n=1) § | DD | RL, RR | 1, 4 | Authors |
| G.CV | Sirsi (n=3) § | EG, MD | RB, RL, RR | 1,1,1 | Authors |
| CV-PG | Wayanad (n=7) §¶ | MD, DD | RL, RR** | 3,4,2 | Authors |
| PG-SG | Parambikulam (n=5)<br>§¶ | MD, DD | RB, RL, RR** | 2,1,3,1 | Authors |
| S.SG | Shenduruney (n=8) §¶ | EG | RB, RL, RR,<br>RR* | 8,1,4,2 | Authors |
| RF data compiled from published literature^ |  |  |  |  |  |
| N.G | Mahabaleshwar⊙ | EG | RR | 6 | Schuller, 1980 |
| G-CV | Kudremukh⊙ | EG | RB, RL, RR | 2,5,6 | Raghuram et al., 2014 |
|  | Moodabidri⊙ | MD | RR | 33 | Chattopadhyay et al.,<br>2010, 2012 |
|  | Srirangapattana⊙ | DD | RR | 6<br>13 | Schuller, 1980;<br>Chattopadhyay et al.,<br>2010, 2012 |
|  | Kodagu ¶ | EG | RB, RL, RR* | n.p. | Villasmil, 2009 |
| PG-SG | Munnar § | MD, EG | RL | n.p. | Sankaran et al., 2012 |
|  | Sirumalai⊙ | EG, MD | RR* | 3 | Chattopadhyay et al.,<br>2012 |
|  | Meghamalai⊙ | EG | RR* | 12 | Chattopadhyay et al.,<br>2010, 2012 |
|  | Valparai ¶ | EG, SHO | RB, RL, RR* | 4,3,8<br>1,2,5<br>2,35,44 | Foui, 2008;<br>Singfield, 2008;<br>Wordley et al., 2014 |
| P.EG | Sivagange §¶ | EG, MD | RB | 2 | Srinivasulu et al.,<br>2014 |
|  | Yercaud ⊙ | EG, MD | RR* | 7 \| 60 | Chattopadhyay et al.,<br>2010, 2012 |
| S.SG | Kalakkad-Mundanthurai<br>Tiger Reserve §¶⊙ | EG, MD | RB, RL, RR* | n.p. | Ponmalar, 2014 |
|  | Kalakkad ⊙ | EG, MD | RR* | 12 | Chattopadhyay, 2010 |
| SL | Haputale, Ratnapura<br>and nearby locations<br>(Sri Lanka) §¶⊙ | MD, DD | RB, RR | 1,1-7,<br>18,13,3,7 | Schnitzler et al., 1985;<br>Neuweiler et al., 1987;<br>Rübsamen and<br>Schafer, 1990;<br>Reimer, 1991;<br>Neumann and<br>Schuller, 1991 |

Key: Biogeographic zones: N.G=North of Goa gap, G-CV=Goa Gap to Cauvery valley, CV-PG=Cauvery valley to Palghat Gap, PG-SG=Palghat Gap to Shenkottah Gap, S.SG=South of
Shenkottah Gap, P.EG=Peninsular India and Eastern Ghats, SL=Sri Lanka; Forest types: DD=Dry-deciduous, MD=Moist-deciduous, EG=Evergreen, SHO=Shola or sub-alpine tropical evergreen forest; Bat species: RR=*R. rouxi*, RR\*=*R. indorouxi*, RR\*\*=both cryptic sibling species co-occur, RB=*R. beddomei*, RL=*R. lepidus*. Symbols provided in the column 'Location' indicate recording conditions: **S** = recorded at roosts, **H** = hand-held recording, **P** = recording from perched bats in flight-cage or captive conditions. Sample sizes provided are numbers of focal bats from which recordings were analyzed: n.p. = not provided in study. Number of roosts or number of call sequences analysed, were often not reported in published studies.

